# Highly reproduceable tapered fiber optoacoustic emitters by in-situ photothermal curing of PDMS for single neuron stimulation

**DOI:** 10.1101/2025.05.24.655947

**Authors:** Guo Chen, Michael Marar, Zhuqin Xu, Wai Yuen Cheng, Feiyuan Yu, Carolyn Marar, Ji-Xin Cheng, Chen Yang

## Abstract

Precise neural modulation is an important tool in neuroscience research for intervene of specific neural pathways. Photoacoustic is an emerging technology enabling high precision non-genetic neural stimulation in vitro and in vivo. In this paper, we report a new method to prepare a Candle Soot - Tapered Fiber based Optoacoustic Emitter (CSTFOE) through an in-situ photothermal curing process. The emitter created was found with high spatial resolution and efficiency for neuromodulation. In the meanwhile, the in-situ photothermal curing procedure is highly reproducible and allows for fabricating the photoacoustic coating on the tiny tip of the tapered fiber with a diameter less than 20 μm. With the high efficiency and reproducibility, we designed an integrated photoacoustic neuromodulation system incorporating the CSTFOE, a nanosecond pulsed laser for photoacoustic generation, and a function generator for synchronization, achieving effective neuromodulation in vitro.

## Introduction

Neurological disorders are the number one cause of disability globally, with cases expected to increase 50% by 2050 (*1*). Therefore, it is critical to continually expand our understanding of neuroscience. The study of neural mechanisms requires devices which have precise neural stimulation capabilities, so even the smallest groups of cells can be monitored in depth.

Among the various neuromodulation techniques, including electrical stimulation (*2*), ultrasound stimulation (*3-16*), infrared stimulation (*17, 18*), optogenetic stimulation (*19*), and photoacoustic stimulation(*20-30*), photoacoustic neuromodulation stands out due to its high precision, non-thermal nature, non-genetic mechanism, and high efficiency. To efficiently generate photoacoustic (PA) wave for neuromodulation, nanosecond pulsed laser was used as the pump beam (*31*). Material with high absorption coefficient and expansion coefficient is preferred as efficient photoacoustic generation material. While PDMS has been well acknowledged as one of the best material as a thermal elastomer with high expansion coefficient (*32*), various different absorbers have been chosen for different purposes. For instance, gold is an effective laser energy absorber and a strong candidate for PA devices. It has been engineered into nanoparticles, rods, spheres, and other nanostructures for PA imaging (*33-36*). However, the narrow absorption band and low stability of gold nanoparticles limit the choice of laser’s excitation wavelength and power (*37*).

Carbon-based materials have emerged as promising alternatives for PA generation (*38*). Graphite-based epoxy has proven viable due to its exceptional laser energy absorption and superior thermal conductivity compared to most other materials (*20*). Carbon nanotubes (CNTs) have also been used to in fiber coatings, demonstrating strong acoustic wave generation (*22*). Among these, candle soot has shown even greater efficiency as a PA converter, owing to its high light absorption and low interfacial thermal resistance (*39*). These properties enable it to generate acoustic waves up to ten times stronger than other carbon-based materials (*23*). Consequently, a candle soot fiber optoacoustic emitter (CSFOE) has been developed to harness this material for precise and effective neuronal modulation (*23*). However, the unique fabrication process involving candle soot particle deposition with PDMS coating presents challenges in coating smaller substrates, such as tapered fibers with a 20 μm diameter. This limitation significantly affects the spatial resolution of stimulation.

In this study, we present a photothermal-induced in-situ coating technique designed for the precise and reproducible fabrication of a PA material—candle soot and PDMS—on the 20 µm tip of a tapered fiber for neuromodulation applications. The fabrication process involves a three-step procedure to create the Candle Soot Tapered Fiber-based Optoacoustic Emitter (CSTFOE), enabling precise control over critical device properties such as layer thickness, absorption characteristics, coating diameter, and PA generation efficiency. With the in-situ photothermal curing process, we demonstrate for the first time that an efficient photoacoustic coating of candle soot/PDMS can be successfully applied to the tiny tip of a tapered optical fiber. This advancement enables both effective photoacoustic neuromodulation and single-cell-level spatial precision simultaneously.

## Results

### Photothermal-induced in-situ curing method for fabrication of CSTFOE

To reproducibly fabricate a photoacoustic coating on the 20 µm tip of a fiber, we developed a three-step method (**Figure 1a)**. The process comprises fiber tapering, candle soot coating, and PDMS photothermal in-situ curing. Bright-field images of the fiber tip after each step are provided in **Figure 1b**.

**Figure 1.**
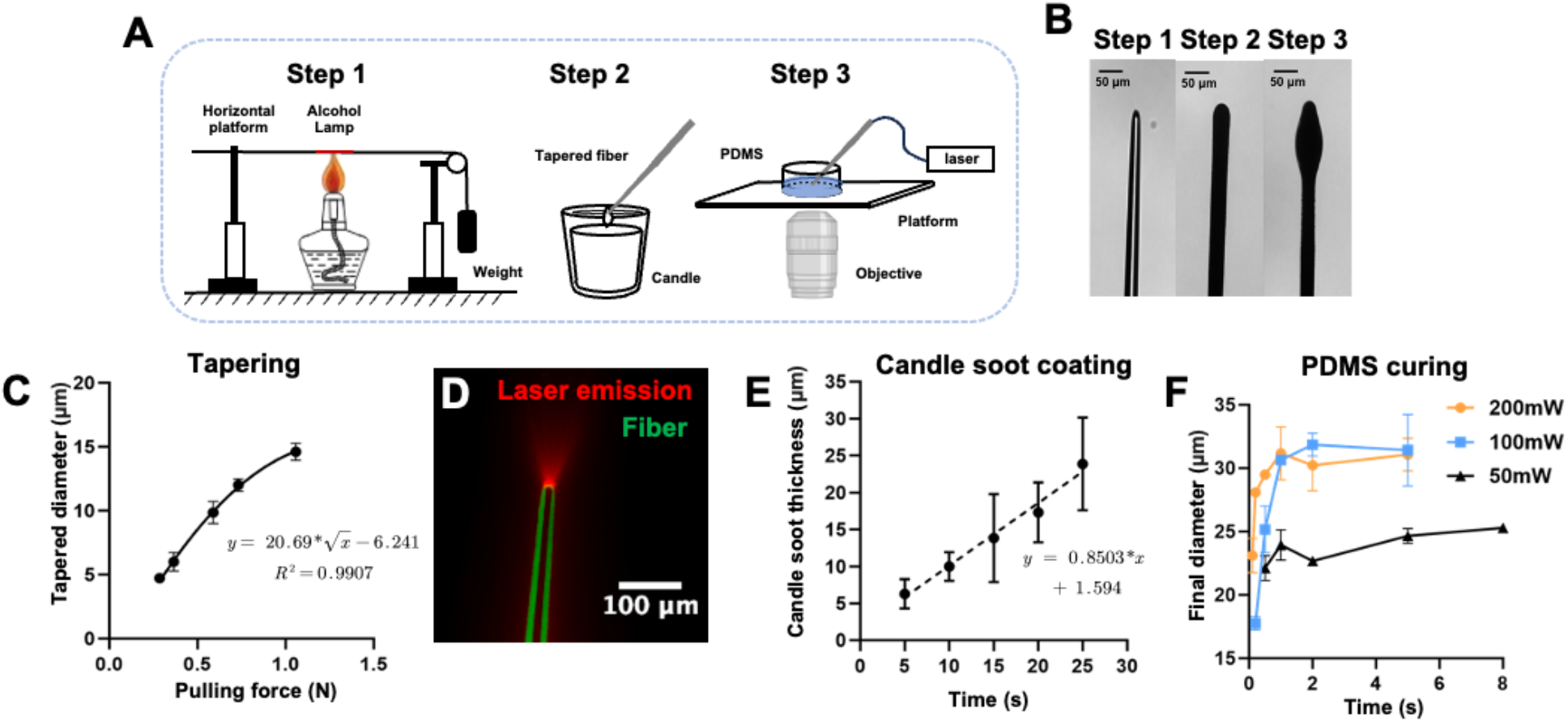
Fabrication and structural characterization of Candle Soot Based Tapered Fiber Optoacoustic Emitter (CSTFOE). A. Schematic of CSTFOE fabrication. B. Pictures after each fabrication step. Scale bar: 50 μm. C. Relationship between the pulling force and tapered diameter. y = 20.69*x^0.5^ – 6.241, R square = 0.9907. N = 42 in total. D. Image for light path of tapered fiber. The fiber is outlined by green color and red is light emitted by fiber. Tip diameter ∼ 15 μm. Laser condition: 1030 nm, cw. E. Relationship between candle soot deposition time and thickness. y = 0.8503*x + 1.594. N = 42 in total. F. Relationship between photothermal curing time and final coating diameter of TFOE in different curing laser powers. N = 23 for 200 mW, N = 17 for 100 mW, N = 10 for 50 mW.

#### Step 1: Fiber Tapering

Fiber tapering is achieved by gradually pulling a 225 µm fiber (FT200EMT, Thorlabs) into a tapered fiber with a tip diameter of less than 20 µm using a thermal tapering method (*22*). The final diameter of the tapered fiber can be tuned by changing the weight of the floating block (**Figure 1c**). The whole tapering process can be modeled and details are in Supplementary Information.

To evaluate the optical performance of the tapered fiber, we measured the laser emission at the tapered fiber tip. First, a bright-field image of the fiber was captured under a microscope (**Figure 1d**, green). Next, the fiber was coupled with a 1064 nm continuous-wave (CW) laser, and the tapered tip was immersed in a scattering medium (5% diluted Intralipid, Sigma) to visualize the emitted laser. The fiber geometry and emitted laser profile were superimposed, as shown in **Figure 1c**. The results confirm that the fiber retains its laser-guiding functionality, emitting light from the tapered tip even after the tapering process.

#### Step 2: Candle soot coating

To deposit a layer of candle soot nanoparticles onto the tip surface of the tapered fiber, the fiber was positioned in the middle of flame of a paraffin wax candle, which is around 2 mm away from the wick. The incomplete burning of the candle produces soot particles (*40*), which adhere to the surface of the tapered fiber. The coating thickness was measured as a function of deposition time, showing a linear relationship (**Figure 1e**). By precisely controlling the coating duration, the thickness of the candle soot layer can be finely tuned with an accuracy of ±5 µm.

#### Step 3: PDMS photothermal in-situ curing

After applying the absorber layer (candle soot), a polydimethylsiloxane (PDMS) layer is introduced as the thermal elastomer to leverage its high thermal expansion coefficient, enabling efficient photoacoustic emission. To coat directly onto the 20 µm tip of the tapered fiber, we developed a photothermal in-situ curing method. To prepare the PDMS, silicone elastomer and curing agent were mixed in a 10:1 ratio. The tapered fiber, with its tip already coated with candle soot, was then immersed in the PDMS mixture. A 1064 nm continuous-wave (CW) laser was coupled into the fiber, delivering the laser to its tip. The candle soot layer absorbed the laser energy, generating localized heat that cured the surrounding PDMS. This process formed a PDMS coating around the fiber tip. The size of the PDMS coating can be changed by varying the laser dosage. With 100 mW laser power and a curing duration of 1 second, the final diameter of the fiber, including the PDMS coating, reached 25 µm (**Figure 1f**). Increasing the laser power resulted in thicker coatings, but the coating thickness plateaued due to reaching thermal equilibrium. This method achieves precise PDMS coating exclusively at the fiber tip, with an accuracy of ±5 µm.

### Characterization of CSTFOE and comparison to CNTTFOE

To quantitatively assess the frequency and spatial distribution of the PA signal emitted by CSTFOE with diameters of 20 μm, the detector’s effective sensing area should be comparable to or smaller than the emitter’s size. However, the smallest commercially available hydrophone has a diameter of 45 μm, which is significantly larger than a typical CSTFOE (20–30 μm), making direct characterization difficult.

To address this limitation, we employed an newly developed optical detection method to measure the PA signal emitted from the CSTFOE [cite SOPPI]. The focused probe beam has a focal diameter of 3 μm, offering significantly improved spatial resolution compared to conventional PA detectors such as hydrophones and transducers (**Figure 2A**). Furthermore, optical detection provides a broad detection bandwidth ranging from sub-MHz to 65 MHz, ensuring more accurate signal measurement than piezoelectric-based PA transducers, which typically have a narrow detection bandwidth (<20 MHz).

**Figure 2.**
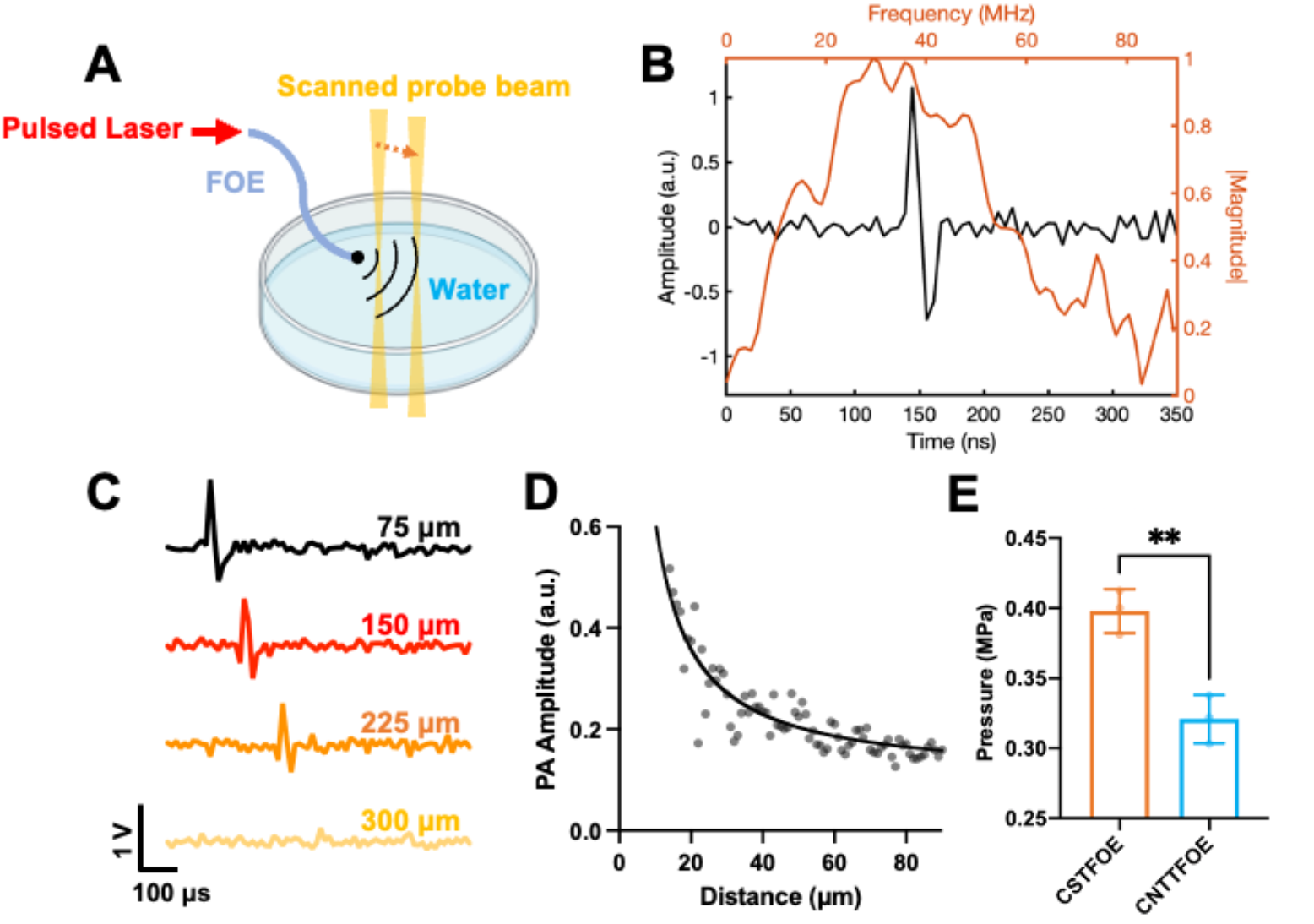
Photoacoustic signal characterization of CSTFOE using optical detection. A. Schematic of photoacoustic measurement using optical detection. B. Representative photoacoustic trace measured from CSTFOE. C. traces of photoacoustic signal detected at different locations plotted as a function of time. D. PA amplitude plotted as a function of distance. E. CSTFOE generates higher pressure than CNT based FOE. N=3 for each group, characterized by hydrophone. *t-test* was performed to calculate the p-value between each group. **: p < 0.01.

The PA signal generated by CSTFOE upon a 1064 nm pulsed laser was detected and plotted as a function of time (**Figure 2B)**, with a central frequency of 30 MHz. By moving the detection beam away from the CSTFOE tip, we measured the PA signals as a function of time (**Figure 2C**), showing amplitude decay with increasing distance. The PA amplitude as a function of distance (**Figure 2D**) exhibits a significant drop within the first 20 μm, demonstrating the high spatial resolution of the PA field emitted by CSTFOE. Moreover, to quantitively compare the pressure generated by CSTFOE and the CNT based TFOE (CNTTFOE) (*22*), we used a hydrophone to measure the PA signals with 1030 nm, 3 ns pulse width and 7.6 μJ pulse energy as pump laser. With its more confined pressure field and the use of candle soot, a higher-efficiency material, CSTFOE generates 1.3 times greater PA pressure than CNTTFOE (**Figure 2E**), highlighting its improved efficiency in both PA generation and spatial precision.

### An integrated system for PA neuromodulation

To enhance the CSTFOE’s compatibility with neuromodulation systems, including calcium imaging and patch-clamp recording, we developed an integrated PA neuromodulation system with all components housed within a compact 12-inch enclosure (**Figure 3A, B**). The system comprises a function generator for laser triggering, a laser control panel, a power supply, and a nanosecond pulsed laser. Additionally, the system features an external synchronization port, enabling seamless integration with recording systems for biological behavior characterization.

**Figure 3.**
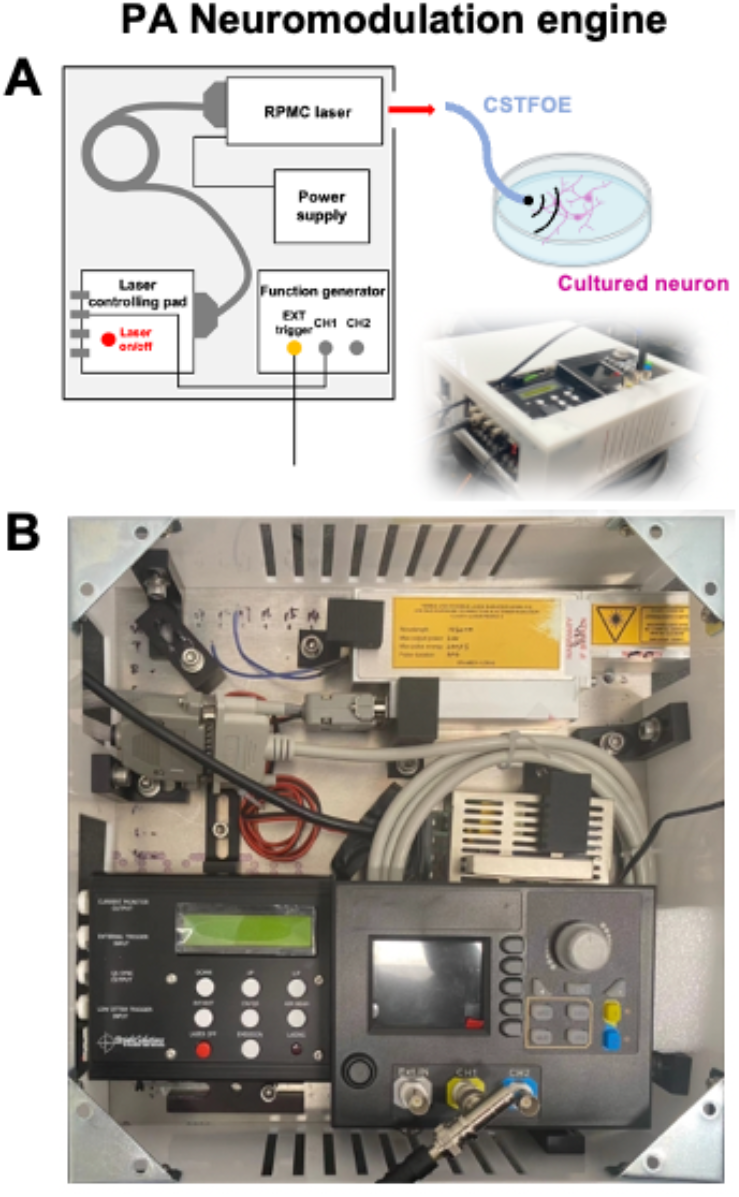
A compact photoacoustic neuromodulation system. A. schematic of PA neuromodulation unit coupled with an CSTFOE for neuromodulation in vitro. B. photo of a neuromodulation engine.

### Neuron stimulation monitored by calcium imaging

With the integrated PA neuromodulation system, the CSTFOE has the ability to stimulate neurons at a sub-100 micron level. A micromanipulator was used to move a CSTFOE within micrometers above a neuron and it was operated under a 1030 nm wavelength laser with 3 ns pulse width at a frequency of 3.3 kHz and a power of 41 mW over a duration of 3 ms. Neural activity was monitored via calcium imaging. Clear stimulation was observed (**Figure 4a**), with an increase in *ΔF/F*_*0*_ of ±193% (**Figure 4b**). To ensure that this level of stimulation was not harmful to the cells, repeated stimulation was performed over the course of two minutes (**Figure 4c**). At 5 s, 35 s, and 65 s the same cell was stimulated, showing a 55-65% increase in *ΔF/F*_*0*_ for all three activations. Therefore, stimulation by the CSTFOE is safe for neurons and shows no deterioration in their performance.

**Figure 4.**
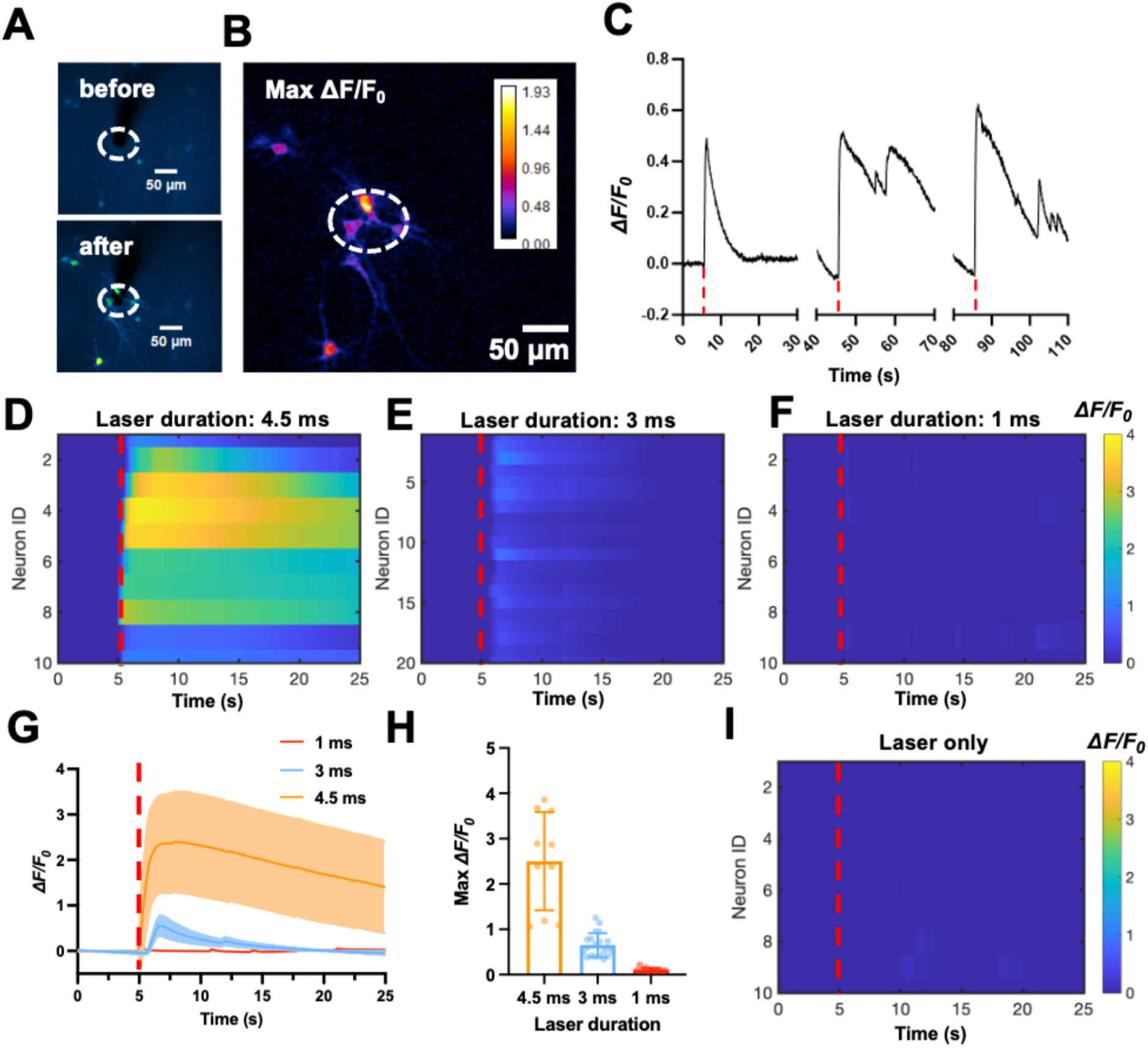
Neuron stimulation monitored by calcium imaging. A. Calcium imaging of GCaMP6f labeled neurons stimulated by CSTFOE before and after stimulation. Scale bar: 50 μm. B. Maximum *ΔF/F*_*0*_ of PA stimulation. C. Calcium trace shows repeatable activation of the same neuron. Laser condition: 3 ms duration, laser power: 41 mW, repetition rate: 3.3 kHz. Red dashed line: laser on. D-F. Colormaps of fluorescence change in neurons stimulated by CSTFOE with different laser burst duration. D. 4.5 ms burst duration. Red dashed line: laser on. E. 3 ms burst duration. F. 1 ms burst duration. G. Average calcium trace traces of neurons attained from (D,E,F) with burst duration of 1 ms (Red) (N = 10), 3 ms (Blue) (N = 20), 4.5 ms (Orange) (N = 10). The shaded region is standard deviation. Laser is on at t = 5 s (red dashed lines). Red dashed line: laser on. H. Average maximum fluorescence intensity changes shown in (D-F). Error bars represent standard deviation. I. Colormap of fluorescence change in neurons around CSTFOE without laser. Red dashed line: laser on.

To examine the effect of different doses of ultrasound on cells, laser duration was varied with constant laser power. Neural activity was monitored via calcium imaging with treatment durations of 4.5 ms, 3 ms, and 1 ms (**Figure. 4d-f)**. A duration of 4.5 ms caused overactivation of neurons, with a 250% increase in *ΔF/F*_*0*_ and an inability for the cells to return to baseline after 20 s afterwards (**Figure 4g-h**). A duration of 3 ms resulted in appropriate neural activation, with an increase of roughly 65%, after which the cells to returned to baseline. A duration of 1 ms showed no significant increase, signifying no activation. To ensure that light leakage was not activating the cells instead of the ultrasound waves, the same experiment was done as the others in this section with a bare tapered fiber (**Figure. 4i**). No activation of the neurons occurred, proving that neuron stimulation is solely due to the PA effect.

## Conclusion

In this paper, we successfully created a candle soot – tapered fiber based optoacoustic emitter (CSTFOE) with a highly quantitative fabrication protocol for neural stimulation at single cell spatial precision. The procedure established to create a CSTFOE allows for variation at any point for a tunable end product, and the simplicity of the procedure leads to a high success rate and reproducibility. Additionally, with higher absorption coefficient of the candle soot layer, this version of an FOE is more efficient than past carbon-based models like the CNT-TFOE, making the CSTFOE a better option for biomedical use.

The primary difference between the CSTFOE and the previous version, being the CSFOE, is the taper of the optical fiber which can create a fiber tip as small as 5 µm before coating. The size of the fiber tip allows for more precise ultrasound waves and easier access to biological environments. With the in-situ photothermal curing in the fabrication process, the PDMS is able to be cured at the tiny tip of the tapered fiber with micron meter level precision.

Characterizing the CSTFOE, a peak-to-peak pressure of about 0.4 MPa is generated, which is larger than the CNTTFOE at 0.32 MPa. Even for a tapered version of these FOEs, it is still evident that candle soot is a more effective absorber for optoacoustic wave generation than CNT. This falls in line with previous observations and confirms that the CSTFOE is the most effective so far for precise neural modulation. Because of the CSTFOE’s capabilities, neuron stimulation was achieved using a laser of 41 mW, 1030 nm wavelength, 3.3 kHz frequency. Experiments showed that at a treatment duration of 3 ms, cell fluorescence increased 55% ± 25, and at a duration of 4.5 ms, fluorescence increased 240% ± 110%. This dosage was shown to be safe for cells.

To summarize, the CSTFOE is able to operate in biological environments efficiently with single cell spatial precision, as proven by the precise stimulation of neurons. Furthermore, the fabrication of the device is customizable and easily reproducible due to its simple three-step procedure. Overall, the CSTFOE provides a neurostimulation platform that can be utilized in neurological research.

### Materials and methods Fabrication of the CSTFOE

**Figure. 1a** shows a schematic of the fabrication process of the CSTFOE. The first step is to create a taper by melting a stock 200 µm optical fiber (FT200EMT, Thorlabs) using an ethanol lamp and then allowing a block of variable weight to pull the two ends of the fiber apart. The second step is to coat the tapered end with candle soot by holding the fiber tip, which is held in a fiber ferrule, inside the flame of a paraffin wax candle for the desired time. The third step is to cure polydimethylsiloxane (PDMS) on top of the candle soot base. The PDMS is made by mixing together silicone elastomer and curing agent in a 10:1 ratio for 5 minutes (Sylgard 184). The CW laser used to thermodynamically cure the PDMS has a 1064 nm wavelength with 400 mW maximum output power (Cobolt Rumba, 1064 nm). Only the tip of the fiber is placed into the uncured PDMS, and it is done precisely using a micromanipulator (MC1000e controller with MX7600R motorized manipulator, Siskiyou Corporation, OR, USA). The fourth step is to use a water sonication machine (Ultrasonic Cleaner, Vevor) to shake off any excess candle soot, and this is done after the fiber has sat at 25°C for 10 hours after the third step so any excess PDMS could cure.

### Characterization of ultrasound waves from optoacoustic effect

For accurate frequency measurements of PA signals from CSTFOEs using optical detection, a continuous wave 1310 nm laser (1310LD-4-0-0, AeroDIODE Corporation) serves as the probe with a power of 5 mW was used to detect the signal. The signal-carrying probe laser was detected by an amplified InGaAs photodiode (PDA05CF2, Thorlabs) with a 1310 nm bandpass filter (FBH1310-12, Thorlabs). The siganl was recorded using a data acquisition card at 180 MSa/s (ATS9462, Alazar Tech), equivalent to 5.6 ns temporal resolution. The signal was then plotted in time trace and frequency spectrum was calculated using Fast Fourier Transform in MATLAB.

For characterizing actually pressure emitted from CSTFOE, a needle hydrophone (HGL-1000, Onda) were used to characterize the signals from the FOEs. The hydrophone was placed into position using a micromanipulator (MC1000e controller with MX7600R motorized manipulator, Siskiyou Corporation, OR, USA), and they were in sub-millimeter range from the tip of the FOEs. The hydrophone’s and FOE’s tips were both completely submerged in water. An oscilloscope (DSO6014A, Agilent Technologies, CA, USA) recorded measurements taken from the hydrophones. A Q-switched 1030 nm laser (Bright Solution, Inc. Calgary Alberta, CA) with a laser pulse of 3 ns was attached to the FOEs to generate PA signal. All PA pressure signals were calibrated from the hydrophones and the frequency data was calculated from a Fast Fourier Transform in MATLAB.

### Neuron culture

Primary cortical neuron cultures were derived from Sprague-Dawley rats. Cortices were dissected from embryonic day 18 (E18) rats of either sex and then digested in papain (0.5 mg/mL in Earle’s balanced salt solution) (Thermo Fisher Scientific Inc., MA). Dissociated cells were washed with and triturated in 10% heat-inactivated fetal bovine serum (FBS, Atlanta Biologicals, GA, USA), 5% heat-inactivated horse serum (HS, Atlanta Biologicals, GA, USA), 2 mM GlutamineDulbecco’s Modifed Eagle Medium (DMEM, Thermo Fisher Scientific Inc., MA, USA), and cultured in cell culture dishes (100 mm diameter) for 30 min at 37°C to eliminate glial cells and fibroblasts. The supernatant containing neurons was collected and seeded on poly-D-lysine coated cover glass and incubated in a humidified atmosphere containing 5% CO2 at 37°C with 10% FBS + 5% HS + 2 mM glutamine DMEM. After 16 h, the medium was replaced with Neurobasal medium (Thermo Fisher Scientific Inc., MA, USA) containing 2% B27 (Thermo Fisher Scientific Inc., MA, USA), 1% N2 (Thermo Fisher Scientific Inc., MA, USA), and 2 mM glutamine (Thermo Fisher Scientific Inc., MA, USA). On day five, cultures were treated with 5 µM FDU (5-fluoro-2’-deox-yuridine, SigmaAldrich, MO, USA) to further reduce the number of glial cells. Additionally on day five, the *pAAV*.*Syn*.*GCaMP6f*.*WPRE*.*SV40* virus (Addgene, MA, USA) was added to the cultures at a final concentration of 1 µL/mL for *GCaMP6f* expression and has cre-independent expression already with no Cre virus involved in the process. Half of the medium was replaced with fresh culture medium every 3–4 days. Cells cultured in vitro for 10–13 days were used for CSFOE stimulation experiment.

### In vitro neuron stimulation

A micromanipulator was used to place the tip of a CSTFOE directly above cells and a Q-switched 1030 nm laser (Bright Solution, Inc. Calgary Alberta, CA) was attached to the CSTFOE. The laser had durations of 4.5 ms, 3 ms, and 1 ms. Calcium fluorescence imaging was performed on a lab-built wide-field fluorescence microscope based on an Olympus IX71 microscope frame with a 20 × air objective (UPLSAPO20X, 0.75NA, Olympus, MA, USA), illuminated by a 470 nm LED (M470L2, Thorlabs, Inc., NJ, USA) and a dichroic mirror (DMLP505R, Thorlabs, Inc., NJ, USA). Image sequences were acquired with a scientific CMOS camera (Zyla 5.5, Andor) at 20 frames per second. Neurons expressing GCaMP6f at DIV (day *in vitro*) 10–13 were used for the stimulation experiment.

### Data analysis

*Prism 10* and *MATLAB* were used to analyze optoacoustic signal data from the hydrophones and to analyze data from the fabrication process of the CSTFOE. Calcium imaging for **Fig. 4a and 4b** were processed using *ImageJ*, and other calcium imaging analysis was done in *Prism 10*. Statistical analysis was completed with *Prism 10* and *Origin 2021*.

