## Supplemental Material for "Highly reproduceable tapered fiber optoacoustic emitters by in-situ photothermal curing of PDMS for single neuron stimulation"

**Physical model of the thermal fiber tapering process**

In the tapering process, the fiber was heated up by an alcohol lamp with a floating block dragging the fiber with a fixed pulling force. When the tapered region of the fiber gets smaller and smaller, the pressure on the cross-section area gets larger and larger, and can be calculated as follows:

$$P = \frac{F}{\pi r^2}$$

Where P is the pressure at the cross-section area in the tapering region, F is the pulling force from the floating block and r is the radium of the tapered region of the fiber. However, at given temperature and material, the ultimate tensile strength (UTS) is determined. When the fiber is tapered thinner and thinner, the pressure increases and when the pressure exceeds the UTS of the fiber, the tapered region will break into two pieces:

$$UTS = P = \frac{F}{\pi r^2}$$

Thus, the tapered fiber's tip diameter can be calculated as below:

$$D = 2\sqrt{\frac{F}{UTS \times \pi}} = \frac{2}{\sqrt{UTS \times \pi}} \times \sqrt{F}$$

From the equation derived above, it can be concluded that the final tip diameter is proportional to the square root of the pulling force, which is consistent with the experiment data. With this physical model, the final tip diameter of the tapered fiber can be precisely controlled by changing the weight of the floating block.
